# Tracing PFAS Transfer from Mother to the Fetoplacental Unit: Insights from Trimester-Specific Maternal Serum Profiles

**DOI:** 10.64898/2026.02.02.703409

**Authors:** Kyle Campbell, Dana Boyd Barr, Andrew J. Morris, Volha Yakimavets, Parinya Panuwet, Donald Turner, Lauren A. Havens, Stephanie M. Eick, Kartik Shankar, Kevin J. Pearson, Aline Andres, Todd M. Everson

## Abstract

Per- and polyfluoroalkyl substances (PFAS) are ubiquitous endocrine-disrupting pollutants that cross the placenta and affect offspring health, but the extent and timing of their transfer to placental and fetal compartments remain poorly understood. We characterized the relationship between trimester-specific prenatal maternal serum PFAS levels and paired placental and cord plasma levels at term. Data came from the Central Arkansas Glowing prospective cohort (n=151, 2010-2014). Four well-detected PFAS were measured using liquid chromatography-tandem mass spectrometry. Regression, elastic net, and parametric g-formula models tested the association between maternal levels in each trimester and placental or cord PFAS levels. In g-formula models, trimester one (T1) or trimester two (T2) measures consistently had the largest effect sizes associated with placental levels (p<0.001-0.05). Similarly, T1 (PFHxS, PFNA, PFOS, PFOA), T2 (PFNA, PFOS, PFOA), and placental PFOA were associated with cord plasma levels (p<0.05-p<0.001). Results were robust to time-varying adjustment for estimated glomerular filtration rate, serum albumin, or maternal weight. Predictive models improved with additional timepoint measures. Our findings suggest PFOA may transfer more efficiently from the placenta to cord plasma and early-to-mid gestation maternal serum PFAS measures may serve as the most robust sentinels of fetoplacental exposure burden, suggesting early exposure prevention should be prioritized.

## INTRODUCTION

Per- and polyfluoroalkyl substances (PFAS) are a large class of persistent organic polyfluorinated compounds with efficient surfactant and repellent properties frequently used industrially and in consumer products worldwide ^1,2^. PFAS burden is widespread in the United States, with detectable levels in essentially all adults, including women of childbearing age^3,4^. PFAS exposure has been associated with a variety of adverse health outcomes^5,6^, including cancer^7^, adiposity^8^, ovarian dysfunction^9,10^, and altered female sex hormones^10^. Prenatal and pediatric PFAS exposure has been associated with abnormal neurodevelopment^11,12^, cardiometabolic outcomes, immune and pubertal dysregulation, and thyroid and renal function^13^. While maternal-to-cord PFAS transfer has been well-documented^14,15^, PFAS dynamics and distributions across maternal, placental, and fetal compartments throughout pregnancy remain unclear.

Epidemiologic investigations of prenatal PFAS have successfully characterized PFAS from critical tissues including maternal blood, the placenta, and umbilical cord blood, suggesting that PFAS that enter maternal circulation can reach the placenta and developing fetus, potentially disrupting the *in utero* environment^14,16–18^. The human placenta is an endocrine organ essential to pregnancy whose function is key to maternal-fetal health^19^. The placenta serves as the gateway to fetal cord blood, a potential barrier against circulating maternal PFAS levels. Recent investigations have attempted to quantify the ability of PFAS to transfer from maternal circulation across the placenta and into cord blood, called the transplacental transfer efficiency (TTE)^15^. TTE is calculated as the ratio of cord blood concentration to maternal concentration of a PFAS congener. However, little is known about how TTE may change throughout pregnancy or how placental PFAS levels may also affect cord blood levels, which represent direct fetal exposure.

We aimed to improve our understanding of the biodistribution of PFAS across pregnancy in a pregnancy cohort from Little Rock and surrounding areas of Central Arkansas. Because PFAS concentrations are highly correlated across repeated maternal serum measures and biologically linked fetoplacental tissues, disentangling the relative contributions of specific exposure windows and biomatrices presents analytic challenges. Thus, we implemented several complementary approaches. First, we estimated maternal serum PFAS transfer efficiency to placental and cord plasma compartments. Second, we examined the individual, joint, and predictive contributions of each repeated maternal serum PFAS measure to placental and cord plasma concentrations. Last, we applied a longitudinal causal inference approach that explicitly accounted for ordered repeated exposures and potential time-varying confounding by pregnancy-related hemodynamic changes. Together, our results provide insight into the biodistribution of PFAS in critical maternal-fetal tissues throughout pregnancy and provide evidence for which biomatrices may best serve as a driver or sentinel of PFAS levels in the placenta or cord blood.

## MATERIALS AND METHODS

### Cohort description

This study was performed in the Glowing Study in Central Arkansas, a prospective birth cohort that recruited pregnant women prior to gestational week 10 between 2010 and 2014 (www.clinicaltrials.gov, ID #NCT03281850)^20,21^. This longitudinal observational cohort consisted of 300 second-parity dyads with singleton pregnancies and mothers who were at least 21 years of age, of whom 153 had successful placenta collection and were eligible for inclusion in this analysis. Glowing enrollment exclusion criteria included self-reported pre-existing medical conditions, sexually transmitted infections, medical complications, reported smoking or alcohol use during pregnancy, medication use known to influence fetal growth during pregnancy, and fertility treatment. Participants self-reported maternal date of birth, maternal education level, child date of birth, and child sex. Maternal age was calculated using self-reported maternal date of birth. All study participants provided informed, written consent and the study was approved by the University of Arkansas for Medical Sciences Institutional Review Board.

### Exposure assessment

Maternal blood samples were collected following an overnight fast in each trimester (trimester 1 >10 weeks ≤ 12 weeks, trimester 2 at 24 weeks, trimester 3 at 30–36 weeks). Umbilical cord blood was collected immediately following delivery. The umbilical cord was doubly clamped after delivery and blood was drawn into 10-mL heparin containing vacutainers. Vacutainers were placed on ice until processed and aliquoted. Both maternal serum and cord plasma were stored at -80°C until PFAS measurement. The PFAS quantification in maternal and cord plasma are detailed elsewhere^22^. Briefly, PFAS were quantified after acetonitrile and methanol treatment to denature protein using ultra-high-performance liquid chromatography-tandem mass spectrometry (UPLC-MS/MS) with electrospray ionization (ESI) using modifications of previously published methods^23–27^. The following PFAS were measured in maternal and cord plasma: Perfluorobutane sulfonate (PFBS), Perfluoropentane sulfonic acid (PFPeS), Perfluorohexane sulfonate (PFHxS), Perfluoroheptanesulfonic acid (PFHpS), Perfluorooctane sulfonic acid (PFOS), Perfluorononanesulfonic acid (PFNS), Perfluorodecanesulfonic acid (PFDS), Perfluoro-n-pentanoic acid (PFPeA), Perfluorohexanoic Acid (PFHxA), Perfluoroheptanoic acid (PFHpA), Perfluorooctanoic acid (PFOA), Perfluorononanoic acid (PFNA), Perfluorodecanoic acid (PFDA), Perfluoroundecanoic acid (PFUnDA), Perfluorododecanoic acid (PFDoA), Perfluorotridecanoic acid (PFTrDA), Perfluorotetradecanoic Acid (PFTeDA), Hexafluoropropylene oxide dimer acid (HFPO-DA; Gen X), 4,8-dioxa-3 H-perfluorononanoic Acid (DONA), 11 -Chloroeicosafluoro-3-oxaundecane-1 -sulfonic acid (11CI-PF30UdS), Perfluorooctanesulfonamide (PFOSA), N-Methylperfluorooctanesulfonamidoacetic acid (N-MePFOSAA), N-ethyl perfluorooctane sulfonamido acetic acid (N-EtFOSAA), 4:2 Fluorotelomer sulfonic acid (4:2 FTS), 6:2 Fluorotelomer sulfonic acid (6:2 FTS), 8:2 Fluorotelomer sulfonic acid (8:2 FTS), and 9-Chlorohexadecafluoro-3-oxanonane-1 -sulfonic acid (9CI-PF30NS). The limit of detection (LOD) for all maternal serum and cord plasma PFAS was 1.0 μg/L.

We have previously discussed placental processing and quantification methods in detail^28^. Briefly, placental core villous tissue was quickly isolated and pooled from both sides and across the placenta and washed of maternal blood after delivery before flash-freezing and pulverization for subsequent PFAS analysis. Placental samples were randomized prior to analysis to reduce analytical batch effects. The 17 target PFAS quantified included: PFHxA, PFHxS, PFHpA, PFOA, PFOS, PFOSA, MePFOSAA, PFNA, PFDA, PFDS, PFUnDA, PFDoA, PFPeA, EtFOSAA, HFPO-DA, PFHpS, and PFBS. The targeted PFAS were analyzed using LC-MS/MS with ESI. The LOD for all placental PFAS was 0.1 ng/g. Concentrations below the LOD were imputed via LOD/√2^29^. Primary models focused on 5 well-detected maternal serum and placental PFAS (PFOS, PFOA, PFHxS, PFNA, and PFDA) and 4 cord PFAS (PFOS, PFOA, PFHxS, and PFNA) with detection rates >55% for each biomatrix (**Supplementary Table 1**).

### Statistical analysis

All statistical analyses were conducted in R v4.3.3^30^. We calculated descriptive statistics of n (%) for categorical variables and median, 1^st^ and 3^rd^ quartiles for continuous variables. We compared descriptive statistics for cohort demographics stratified by participants that were included vs. excluded with a Chi-squared test for categorical variables and the Wilcoxon rank sum test for numeric variables. We calculated Spearman correlations between PFAS measures. Each model tested a single pollutant, was case-complete, and adjusted for maternal age and education, baseline body mass index (trimester one), and fetal sex with placental (n=99) or cord (n=67) PFAS levels as the outcome. Adjustment variables were selected a priori from a causal inference perspective with directed acyclic graphs (DAGs). Each PFAS predictor was log_2_-transformed to better approximate a normal distribution for modelling purposes. We used mixed effects linear models to test log-transformed transplacental transfer efficiency (TTE) changes over pregnancy with a random intercept.

Because PFAS concentrations were highly correlated across repeated maternal measures and biologically linked tissues, we implemented a complementary analytic framework designed to address distinct inferential and prediction goals (Table 1). Conventional linear regression models were used for simple association testing and comparability with prior literature. Penalized regression approaches (LASSO, elastic net, and ridge regression) were then used to evaluate relative variable importance while accounting for multicollinearity among PFAS measures across time and tissues. To explicitly model the ordered, time-varying structure of our PFAS measurements, we applied parametric g-formula within the stochastic intervention framework. Statistical significance was assessed at alpha = 0.05. Finally, because predictors that optimize exposure prediction may differ from those that yield interpretable inferential estimates, we evaluated predictive performance using cross-validated linear prediction models.

**Table 1.**
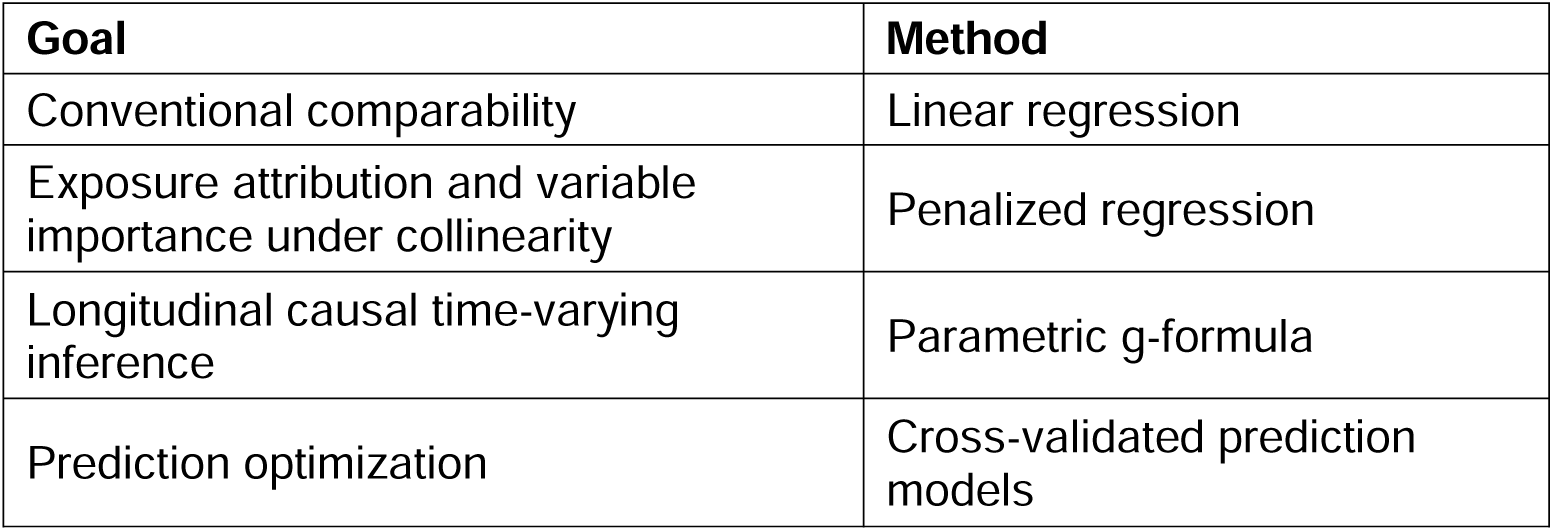
Goals and analytic approaches utilized to achieve those goals.

We first evaluated whether maternal trimester-specific PFAS concentrations predicted placental PFAS levels, and whether maternal and placental PFAS concentrations predicted cord plasma PFAS levels using simple and multivariable linear regression models. We next applied penalized regression approaches to estimate relative variable importance while reducing instability from multicollinearity. Shrinkage and regularization models included LASSO (alpha = 1), Elastic net (alpha = 0.5), and Ridge regression (alpha = 0) models implemented in glmnet^31^, R package v4.1-8. Penalty parameters (λ) were selected through cross-validation and adjustment covariates were forced into the model.

To evaluate exposure prediction independent of inferential interpretation, we applied 10-fold cross-validated predictive linear models to identify combinations of biomatrices that optimized prediction accuracy for placental and cord plasma PFAS concentrations. Predictive performance was assessed using root mean square error (RMSE) implemented in xvalglms, R package v.0.1.8^32^.

To explicitly model the ordered time-varying structure of PFAS concentrations across pregnancy, we applied the parametric g-formula^33^ through a stochastic intervention regime^34^ framework. With this approach, we estimated changes in placental and cord plasma PFAS concentrations under hypothetical shifts in maternal PFAS concentration distributions at each trimester. We adapted a recently published approach for identifying influential mixture components^35^, and estimated the effect of increasing each PFAS concentration by 25% of its pooled standard deviation relative to naturally observed values; maternal serum PFAS measures were pooled across timepoints to calculate their standard deviations. Models were implemented in gfoRmula^36^, R package v1.1.1, with placental or cord plasma PFAS concentrations as continuous outcomes. Regression models were fit for each time-varying exposure, time-varying covariate, mediator, and outcome, conditional on prior exposure and covariate history at each time point. The fitted models simulated longitudinal data under specified interventions with Monte Carlo simulation using 500 bootstrap samples.

We conducted three sensitivity analyses to evaluate potential time-varying confounding by maternal serum albumin, estimated glomerular filtration rate (eGFR), or maternal weight measured at each trimester. DAGs were utilized to illustrate modelling assumptions and putative relationships among exposures, mediators, covariates, and outcomes (**Supplementary Figures 1-2**).

## RESULTS AND DISCUSSION

### Descriptive statistics

Of 153 mother-child dyads with placental samples, two samples had insufficient quantity of tissue to perform quantification. One individual was additionally excluded due to missing covariate information. This study focused on the Glowing participants who were case-complete for maternal serum across all three trimesters and for placental PFAS quantification (n=99), or who were case-complete for maternal serum across all three trimesters, placental and cord plasma PFAS quantification (n=67). Concentrations of four PFAS (PFOS, PFOA, PFNA, and PFHxS) were detectable in over 55% of placental, maternal serum, and cord plasma samples and were the primary focus of downstream analyses. PFDA was also analyzed in models of placental PFAS, since it met this cutoff for placenta but not for cord plasma levels. Both case-complete samples were sex-balanced, had median gestational ages of 39.3 weeks, median maternal ages of around 30 years old, and about a third of each sample had a college education or greater (**Table 2**). There were no statistically significant differences (p<0.05) in demographic covariates or PFAS measures in the placental nor cord outcome samples between included and excluded samples, except for a modest difference (p=0.021) in placental PFOS concentrations in the placental outcome dataset based on a median concentration of 0.39 ng/g in the included sample compared to 0.48 ng/g in the excluded sample (**Supplementary Tables 2-3**). PFAS measures exhibited strong Spearman correlations (p<0.05) across biomatrices and mostly correlated across PFAS within tissue (**Supplementary Figure 3**).

**Table 2.** Descriptive statistics of the analytic datasets modelling either placental PFAS concentrations as the outcome or.

|  | Placenta outcome | Cord blood outcome |
| --- | --- | --- |
| Characteristic | N = 99<sup>1</sup> | N = 67<sup>1</sup> |
| Child sex |  |  |
| Female | 45 (45%) | 33 (49%) |
| Male | 54 (55%) | 34 (51%) |
| Characteristic | N = 99<sup>1</sup> | N = 67<sup>1</sup> |
| Gestational age (wks) | 39.29 (39.00, 39.71) | 39.29 (39.00, 39.86) |
| Maternal age | 29.90 (27.90, 32.49) | 29.67 (28.25, 32.88) |
| Maternal education |  |  |
| Less than college | 33 (33%) | 23 (34%) |
| College or greater | 66 (67%) | 44 (66%) |
| Maternal race |  |  |
| African American | 9 (9.1%) | 3 (4.5%) |
| Asian | 1 (1.0%) | 1 (1.5%) |
| More than One Race | 1 (1.0%) | 0 (0.0%) |
| **Characteristic** | **N = 99<sup>1</sup>** | **N = 67<sup>1</sup>** |
| Unknown or Not Reported | 1 (1.0%) | 1 (1.5%) |
| White | 87 (88%) | 62 (93%) |
| Maternal ethnicity |  |  |
| Hispanic | 3 (3.0%) | 2 (3.0%) |
| Non-Hispanic | 95 (96%) | 64 (96%) |
| Unknown | 1 (1.0%) | 1 (1.5%) |
| Baseline BMI | 25.1 (22.2, 28.7) | 24.6 (22.0, 28.3) |

<sup>1</sup> n (%); Median (Q1, Q3); Glowing, Central Arkansas, 2010-2014

### Transplacental transfer efficiency increased throughout pregnancy

To identify potential differences in TTE for each PFAS and trimester, we calculated trimester-specific and PFAS-specific TTEs (**Figure 1A**) and modeled their association with gestation progression. Geometric means for PFAS-specific TTEs ranged from 27.9% for PFOS in trimester one to 72.1% for PFNA in trimester three (**Table 3**). For each successive trimester, TTE was associated with a 1.14x increase for PFOS (p<0.001) and 1.29x increase for PFNA (p<0.001). We then calculated placental transfer ratios to measure relative PFAS transfer across maternal serum to placenta and placenta to cord plasma^37^ by dividing placental concentrations by serum or plasma concentrations and multiplying by the average human serum density of 1.018 g/mL^38^ or the average human plasma density of 1.025 g/mL^39^ (**Figure 1B**). Overall, geometric means for PFAS-specific TTEs tended to be lowest for PFOS, while placental transfer ratios were lowest for PFOA across all trimesters, and trimester 3 had the largest TTEs and placental transfer ratios for all PFAS (**Table 3**). We found no associations between potential hemodynamic-related measures maternal serum albumin or eGFR and any TTEs (data not shown).

**Figure 1.**
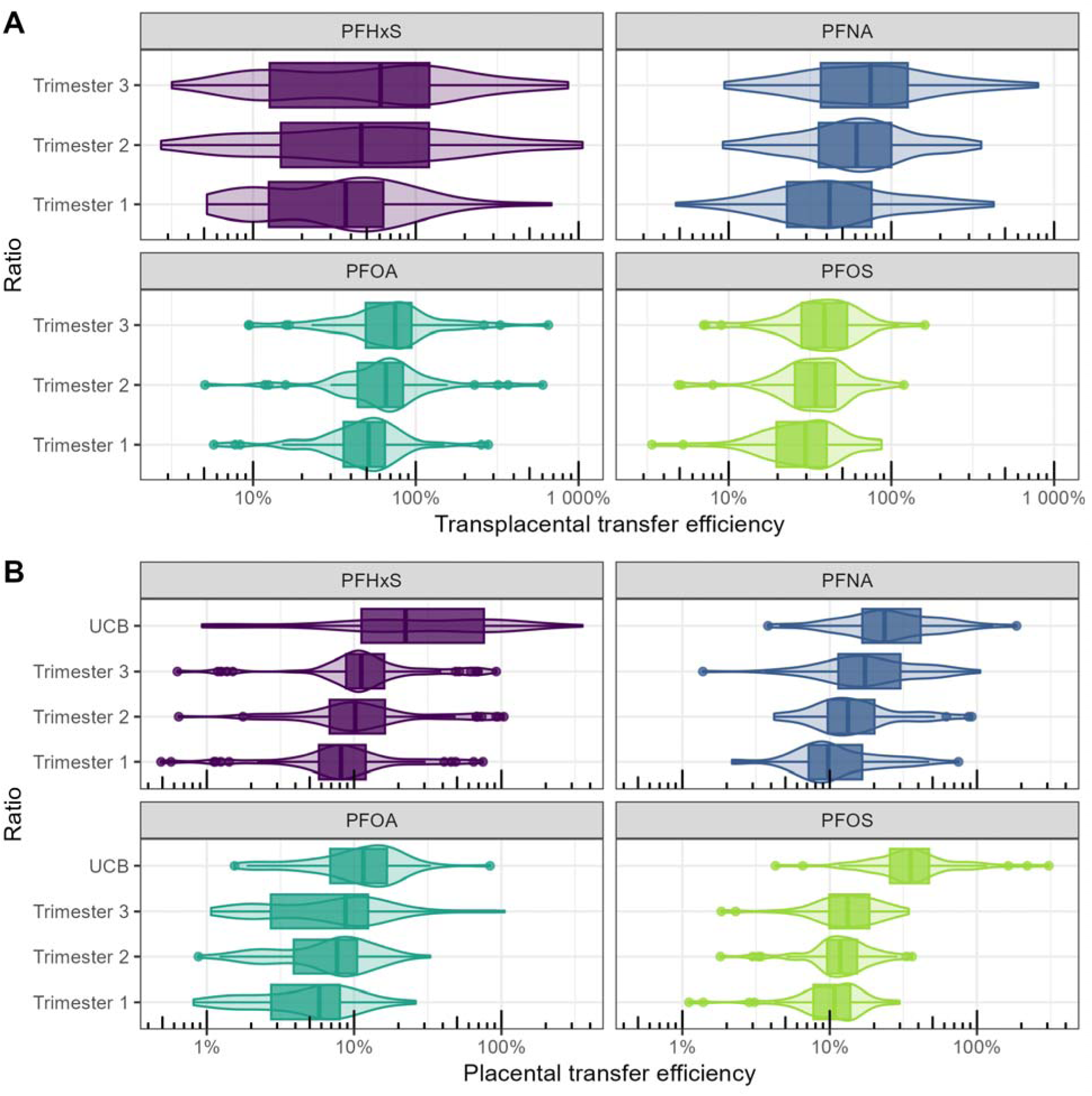
**(A)** Transplacental transfer efficiency for each PFAS at each timepoint. **(B)** Placental transfer efficiency for each PFAS at each timepoint. Transfer efficiency ratios are expressed as percentages, where 100% indicates equivalent concentrations between compartments

**Table 3.** Geometric means for PFAS-specific transplacental transfer efficiencies (TTEs) and placental transfer ratios alongside mixed model results estimating the fold-change in TTE per trimester for each PFAS. TTE means are expressed as the percentage of cord plasma PFAS concentration divided by maternal serum concentration. Placental transfer ratio means are expressed as a percentage by dividing placental PFAS concentrations by maternal serum concentrations and multiplying by the average human serum density of 1.018 g/mL or by cord plasma concentrations and multiplying by the average human plasma serum density of 1.025 g/mL for UCB-Placental.

| *Geometric mean of transfer efficiency (%)* | *Trimester 1 TTE* | *Trimester 2 TTE* | *Trimester 3 TTE* | *Trimester 1- Placental* | *Trimester 2- Placental* | *Trimester 3- Placental* | *UCB- Placental* | *Transplacental transfer efficiency mixed model fold-change (per trimester)* |
| --- | --- | --- | --- | --- | --- | --- | --- | --- |
| *PFNA (C9)* | 43.4 | 59.2 | 72.1 | 10.8 | 14.7 | 18.0 | 25.1 | 1.29 (1.17, 1.43) |
| *PFOA (C8)* | 46.6 | 63.6 | 66.9 | 4.7 | 6.5 | 6.8 | 10.2 | 1.20 (1.09, 1.31) |
| *PFOS (S8)* | 27.9 | 32.8 | 36.1 | 9.9 | 11.6 | 12.8 | 35.6 | 1.14 (1.09, 1.20) |
| *PFHxS (S6)* | 33.4 | 46.8 | 49.5 | 7.9 | 11.1 | 11.7 | 23.8 | 1.22 (1.08, 1.39) |

### Trimester-specific maternal PFAS associations with placental PFAS measures varied by PFAS

We next tested which trimester-specific maternal PFAS measurements were associated with placental PFAS levels at term using both individually modeled trimester-specific regressions and jointly adjusted models that included all three maternal timepoints simultaneously (**Figure 2, Supplementary Table 4**). All linear PFAS trimester-specific measures were associated with their respective placental PFAS concentrations in single timepoint models, and effect estimates were largely consistent across trimesters. In the co-adjusted GLM models, first trimester coefficients were attenuated to the null for most PFAS: PFOS (p=0.31), PFOA (p=0.29), and PFHxS (p=0.17). Second trimester PFDA (p=0.06) and third trimester PFNA (p=0.13) were also pulled to the null with co-adjustment. These results were consistent with LASSO, elastic net, and ridge regularization models, with point estimates largely comparable to the joint GLM estimates. Placental PFAS levels were most strongly informed by second and third trimester PFOS and PFHxS concentrations, second trimester PFOA, and first and second trimester PFNA levels, whereas PFDA concentrations across pregnancy contributed relatively equally across all three trimesters.

**Figure 2.**
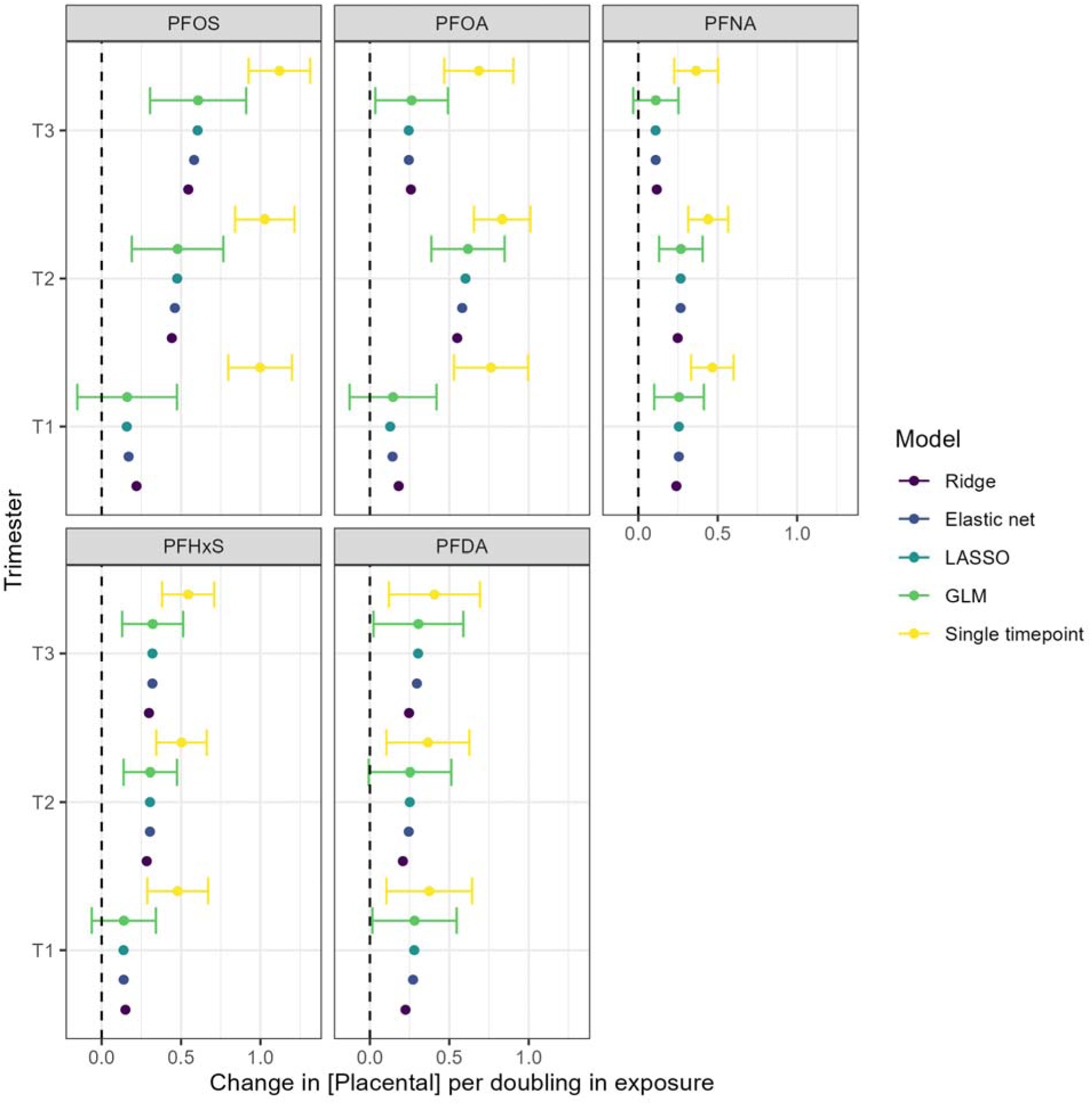
Linear model results of the association between a doubling in each trimester PFAS measure (μg/L) individually and placental PFAS concentration (ng/g) at a single timepoint and jointly, including regularized regression models (Elastic net, ridge, and LASSO), which do not provide precision estimates.

### Stochastic intervention models identify early trimesters as primary drivers of placental PFAS levels

Using the stochastic intervention framework, we estimated the relative time-ordered contribution of trimester-specific maternal PFAS concentrations to term placental PFAS levels by modelling the expected change in placental PFAS associated with a 25% standard deviation increase in maternal PFAS concentrations at each trimester, while holding other exposures at their naturally observed levels. Stochastic interventions at all timepoints for all PFAS were associated with increased placental PFAS burden (p<0.05, **Figure 3, Supplementary Table 5**). PFDA, the least well-detected PFAS, had similar effect estimates regardless of the intervention timing. For all other PFAS, first trimester (PFHxS, PFNA, PFOS) or second trimester (PFOA) was the largest driver of placental PFAS levels. These findings were largely consonant with the relative importance rankings in the conventional and regularization-based models for PFDA and PFOA. In contrast, the g-formula-based estimates favored the first trimester followed closely by the second trimester as the largest contributor of placental PFAS levels for PFHxS, PFNA, and PFOS.

**Figure 3.**
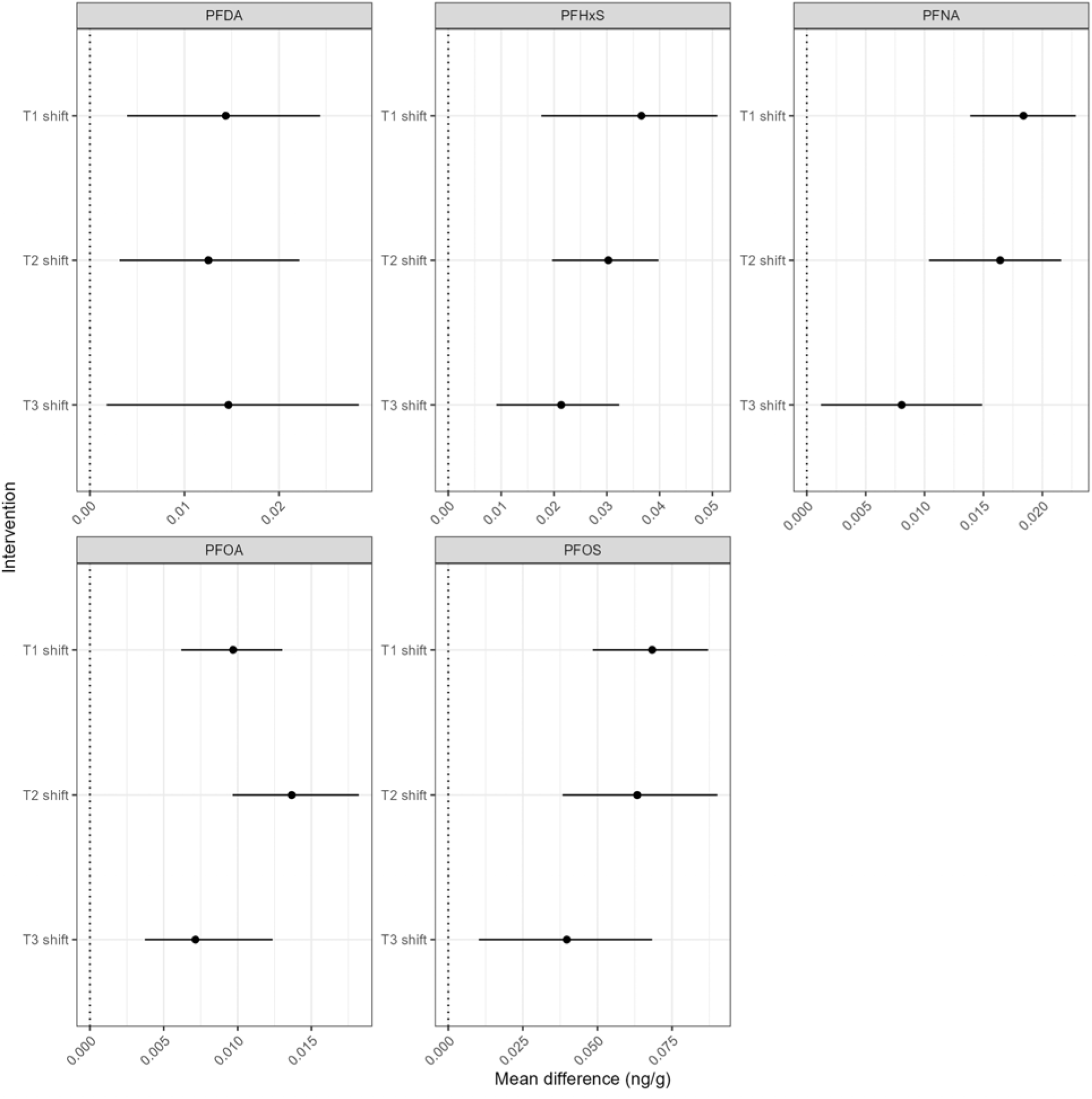
Stochastic intervention estimates of the mean difference in placental concentration corresponding to a 25% standard deviation increase (shift) at one timepoint during pregnancy via the parametric g-formula.

Sensitivity analyses additionally evaluated potential time-varying confounding by maternal serum albumin, eGFR, and maternal weight across pregnancy. Three individuals were additionally excluded for not having complete hemodynamic measures at each timepoint and one was excluded for not having complete weight information available at each timepoint, leaving n=95 individuals for these sensitivity analyses and the primary stochastic intervention analysis for comparability. Effect estimates were consistent across models, suggesting minimal influence of these physiological and body composition factors on the observed associations (**Supplementary Figure 4**).

### Predictive modelling of placental PFAS measures improved by integrating across multiple measures

Finally, to identify the best predictor of placental PFAS measures, we used 10-fold cross-validated linear models with 200 repeats using all combinations of maternal trimester-specific PFAS measures adjusted for covariates, with root mean square error of prediction (RMSE_P) to evaluate predictive performance (**Figure 4**). Across all five PFAS, the models that included second and third trimesters or all three trimesters resulted in the lowest RMSE_P. However, when only considering a single timepoint, second trimester PFOA stood out as the biggest contributor to performance for PFOA, as well as the first and second trimesters for PFNA for placenta PFNA. These results suggest that mid-gestation maternal serum measures may serve as better biomarkers for placental PFOA or PFNA. For PFOS and PFHxS, the third trimester was the single most important timepoint for improving prediction. Overall, while some trimester-specific patterns emerged, incorporating more measures across gestation generally provided more accurate prediction of placental PFAS levels.

**Figure 4.**
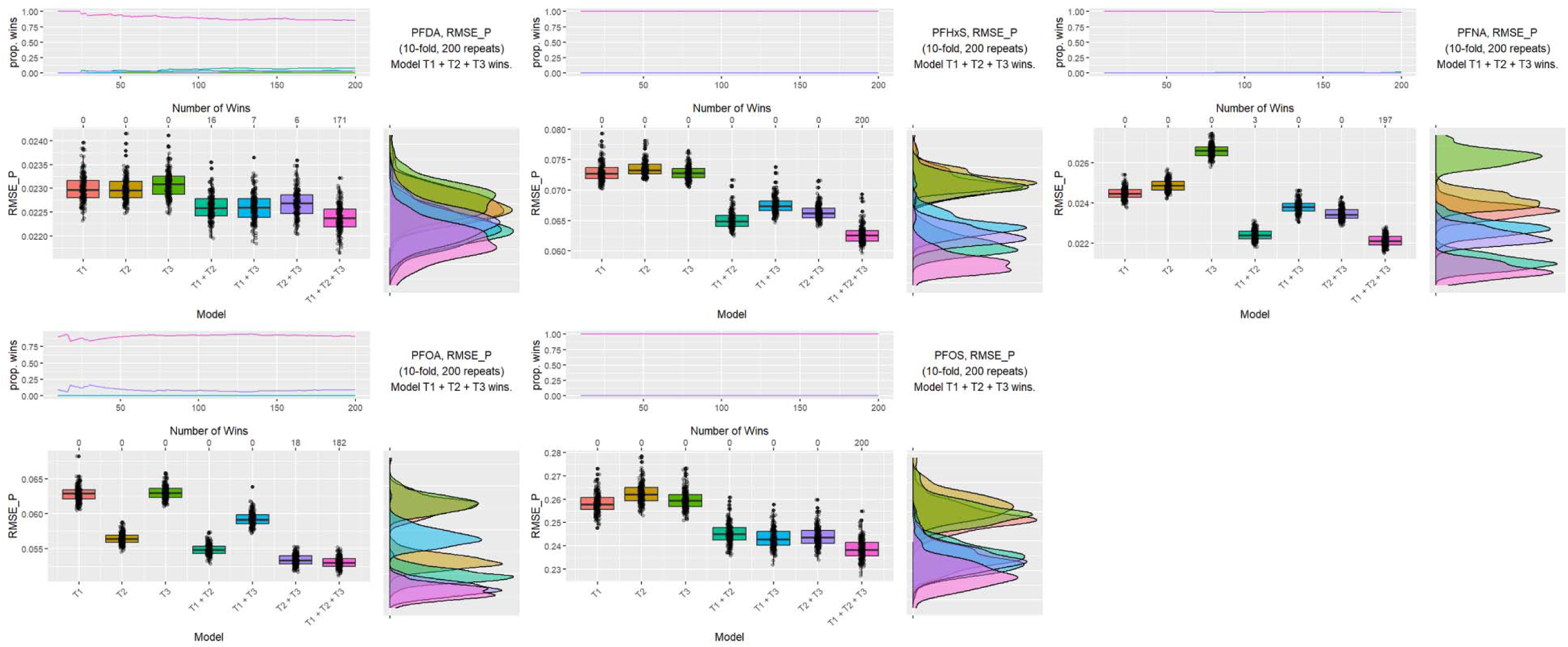
Cross-validated linear models were used to identify which combination of maternal PFAS measures that yield the lowest RMSE_P. Four panels are presented for each PFAS (PFDA, PFHxS, PFNA, PFOA, and PFOS), with the top left panel representing the proportion of iterations in which that biomatrix combination was selected as the best prediction as folds progressed, with colors corresponding to the box plots of the RMSE_P in the bottom left panel and the density plots of RMSE_P on the bottom right panel. The most predictive combination of biomatrices for each placental PFAS is presented in the top right panel.

### Trimester-specific maternal and placental PFAS associations with cord plasma PFAS measures varied by PFAS

To identify which trimester-specific maternal serum PFAS measures and placental PFAS at term were associated with cord plasma PFAS measures, we applied the same framework as above. In conventional regression models, PFOS, PFOA, and PFNA levels in all biomatrices were significantly positively associated with cord plasma levels, whereas only trimester one PFHxS was associated with cord plasma levels (**Figure 5, Supplementary Table 6**). Co-adjustment again attenuated most associations and revealed substantial variability in the biomatrices that were most informative of cord plasma PFAS concentrations. Placental PFOA, second-trimester PFNA, and first-trimester PFHxS were the strongest predictors of corresponding cord plasma levels, whereas maternal and placental PFOS measures were similarly informative of cord plasma PFOS concentrations.

**Figure 5.**
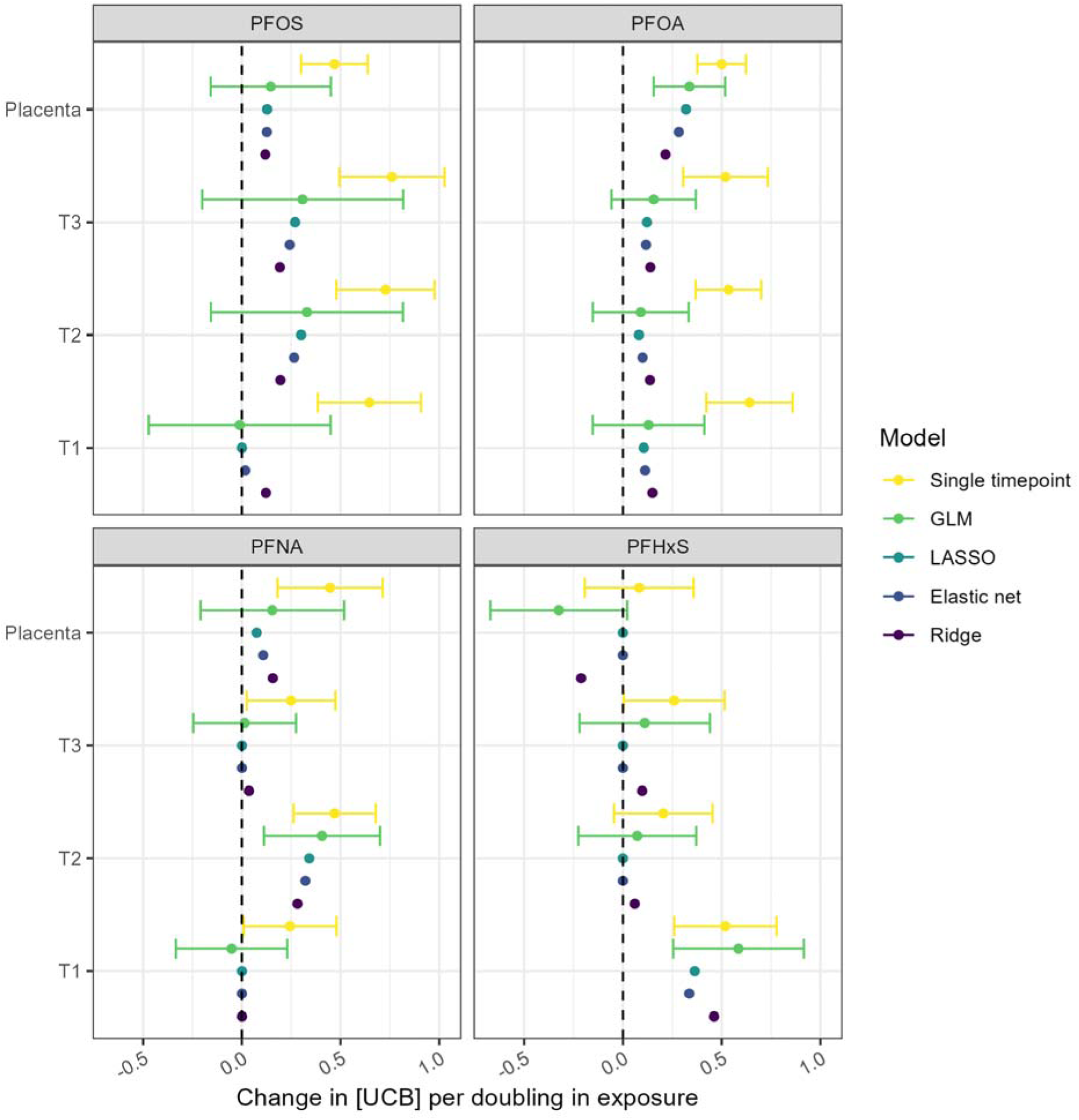
(**A**) Linear model results for the association between a doubling in each trimester PFAS measure (μg/L) and cord blood serum PFAS concentration (μg/L) individually at a single timepoint and jointly, including regularized regression models, which do not provide precision measures.

### Stochastic intervention models identify early trimesters as primary drivers of cord plasma PFAS levels

The stochastic intervention regime showed that first-trimester PFHxS, second-trimester PFNA, and both early/mid-pregnancy PFOS measures showed the strongest estimated contributions to corresponding cord plasma levels, whereas PFOA contributions were more broadly distributed across pregnancy and placental compartments, with no single biomatrix clearly dominating prediction (**Figure 6, Supplementary Table 7**). Similar to the placenta outcome models, there was no evidence of time-varying confounding by albumin, eGFR, or maternal weight in g-formula sensitivity analyses of the cord plasma outcome models (**Supplementary Figure 5**); two individuals were additionally excluded for not having complete hemodynamic and weight data at every timepoint, leaving n=65 individuals for these sensitivity analyses and the primary stochastic intervention analysis for comparability. Overall, early trimester biomatrices were most informative for PFOS, PFHxS, PFNA, and PFOA while placental levels were additionally and uniquely informative for cord plasma PFOA.

**Figure 6.**
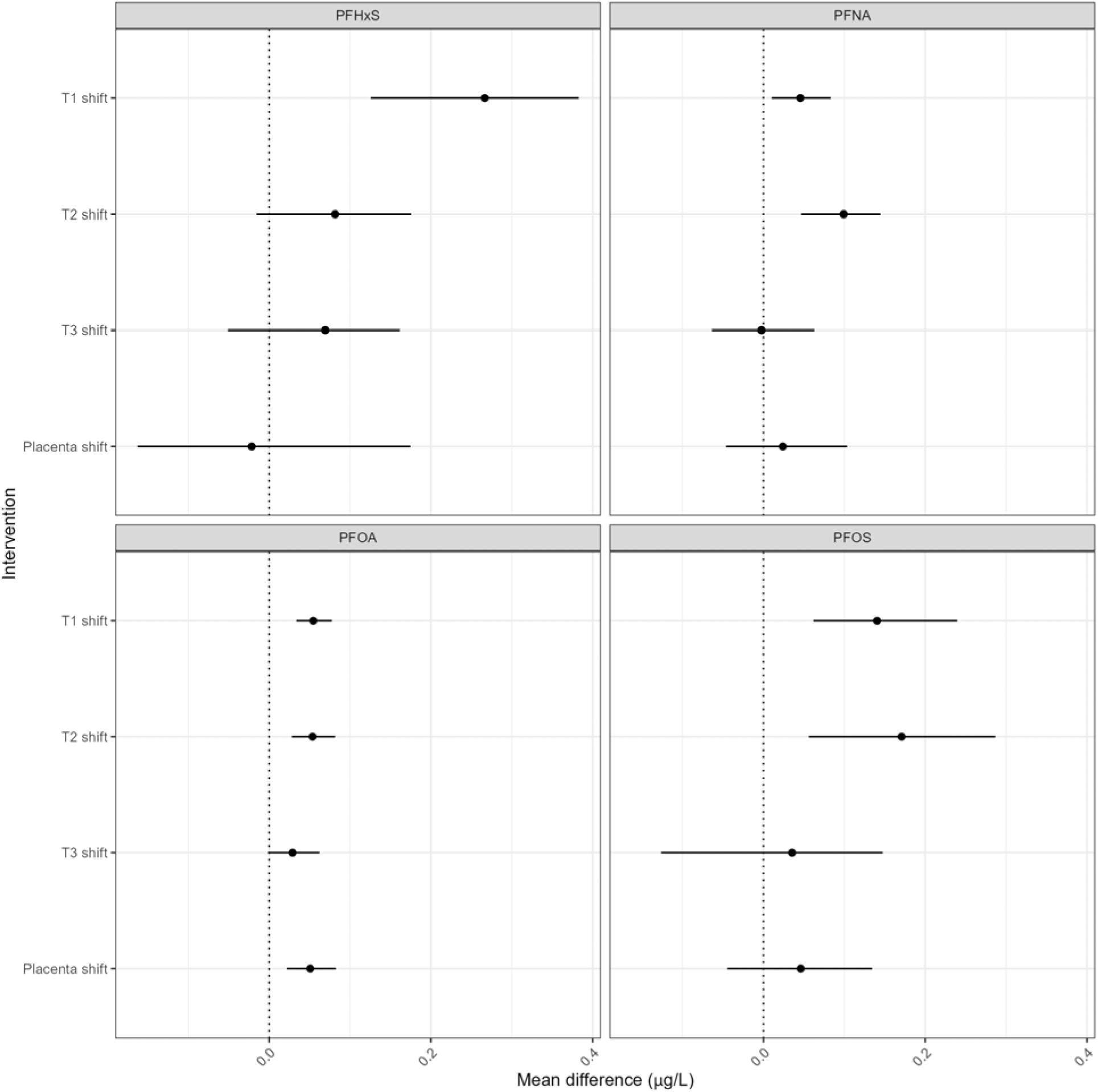
Stochastic intervention estimates of the mean difference in cord plasma concentration corresponding to a 25% standard deviation increase (shift) at one timepoint during pregnancy via the parametric g-formula.

### Predictive models of cord plasma PFAS measures varied by PFAS species, but early and mid-pregnancy serum levels tended to be most informative

To identify the best predictor of cord plasma PFAS measures, we applied the same cross-validation framework as before (**Figure 7**). Among the single-sample prediction models, the second trimester provided the best prediction for PFOS and PFNA, while placenta was the best predictor for PFOA and the first trimester was the best predictor for PFHxS. For PFOS, PFOA and PFHxS, integrating information from more than one biomatrix improved prediction of cord plasma levels with the second plus third trimesters being the best overall predictor for PFOS, and with the first trimester plus placenta being the best overall predictor for PFOA and PFHxS. For PFNA, the second trimester alone was the best overall prediction model, even better than models that incorporated additional information from other samples. These findings largely align with the above modelling results and underscore the predictive performance of early-and mid-pregnancy serum PFAS measures for predicting cord plasma levels at term, as well as placenta being particularly informative for PFOA cord plasma levels.

**Figure 7.**
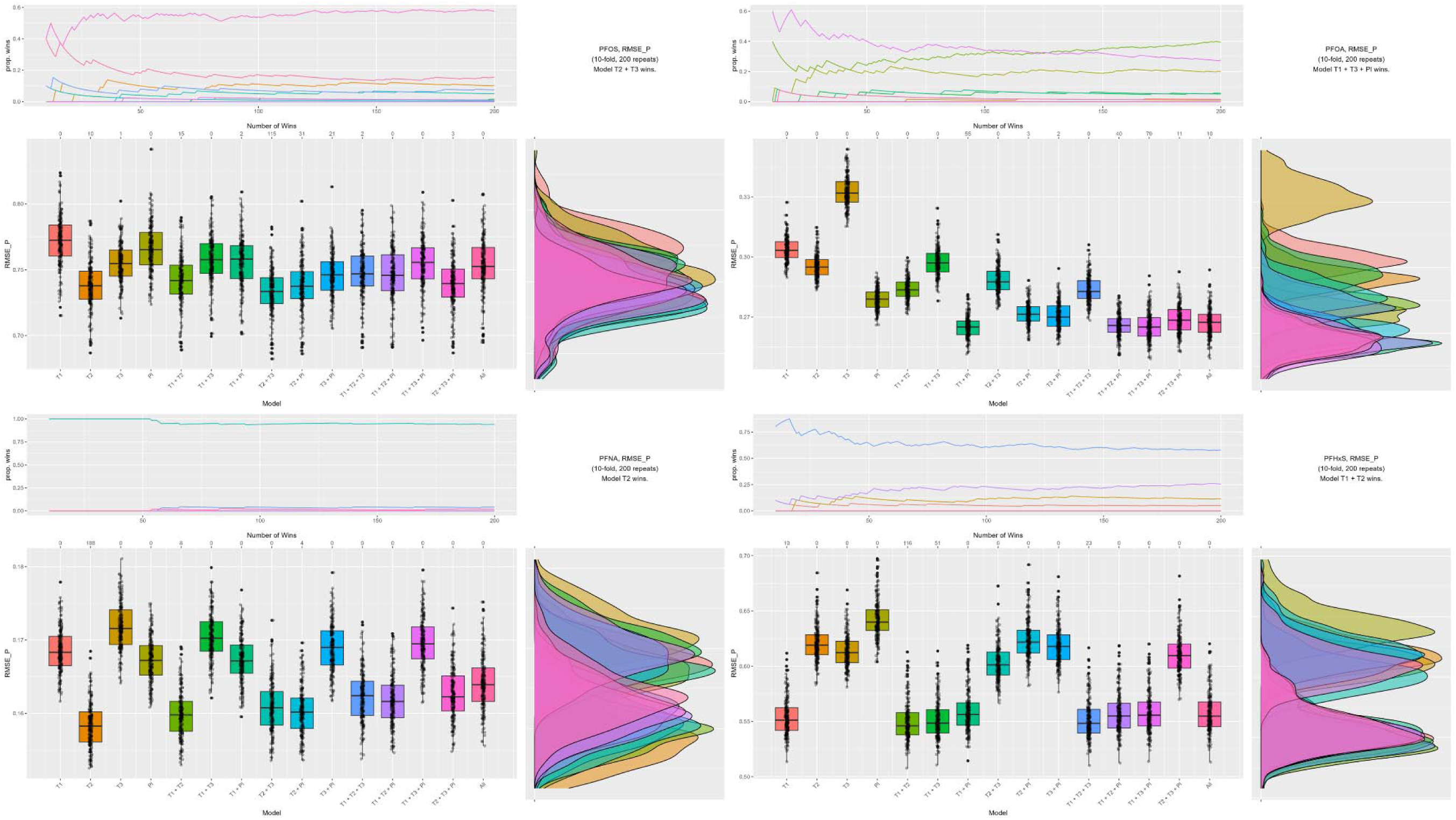
Cross-validated linear models were used to identify which combination of maternal and placental PFAS measures that yield the lowest RMSE_P for predicting cord plasma levels. Four panels are presented for each PFAS (PFOS, PFOA, PFNA, and PFHxS), with the top left panel representing the proportion of iterations in which that biomatrix combination was selected as the best prediction as folds progressed, with colors corresponding to the box plots of the RMSE_P in the bottom left panel and the density plots of RMSE_P on the bottom right panel. The most predictive combination of biomatrices for each placental PFAS is presented in the top right panel.

We leveraged the unique Glowing pregnancy cohort, which collected one of the most comprehensive sets of repeated PFAS measurements, including longitudinal maternal serum, as well as placental, and cord plasma samples to test how prenatal exposures influence fetoplacental PFAS burden. Together, our results suggest that early-to-mid, especially second trimester PFAS measures from maternal serum, were predominately the most important sentinels of placental and cord plasma levels for almost all PFAS. Nonetheless, maternal serum at each timepoint meaningfully contributed to placental levels at delivery. First trimester was the most critical timepoint for PFHxS for determining placental or cord plasma levels. Placental PFOA was also an important predictor of PFOA cord plasma levels. For accurately predicting placental or cord plasma PFAS levels, incorporating multiple measures at these timepoints and biomatrices reduced prediction error.

Our findings were consistent with those of previously published studies in several significant ways. First, PFOS and PFOA were the most abundantly detected PFAS in our sample across all timepoints and tissues. As previously reported by our group, maternal levels in Glowing for PFOS and PFHxS are comparable to NHANES 2011-2014 levels whereas PFNA and PFOA were elevated^22^. Glowing maternal PFAS levels were similar, but slightly lower than more recent NHANES 2017-March 2020 pre-pandemic data.^4^ Glowing placental PFOS and PFOA levels were similar to a Durham, North Carolina-based cohort^16^, but lower than several cross-sectional studies from Denmark^40,41^ and a China-based cohort study^42^. We observed a downward trend of TTE as gestation progressed for each PFAS tested, which corresponds to a decreasing concentration of PFAS throughout gestation, a trend that has been previously observed^43,44^, but inconsistently in different populations, such as an Atlanta-based African American population where the trend was reversed^14,15^. Finally, PFAS with sulfonyl groups, PFOS and PFHxS, had the lowest TTEs across gestation, consistent with a large meta-analysis of TTEs^15^.

The repeated maternal serum measures in Glowing provided the opportunity to calculate TTE at multiple timepoints in the same individuals with the same approach. Trends in TTE by carbon chain length and functional group were consistent with prior findings, except for PFNA where absolute TTE values were typically lower than those previously reported for earlier gestational timepoints. The heterogeneity in TTE effect estimates across pregnancy in this cohort could highlight the importance of timing when measuring PFAS from maternal biomatrices during pregnancy, a previously reported concern^15,44^, that could have effects on estimated TTE and our interpretation of PFAS transfer within and between the fetal and maternal compartments. One potential explanation for the change in PFAS levels and corresponding changes in estimated TTEs throughout gestation is the changes in blood volume that occur as pregnancy progresses. We attempted to replicate a previously identified association between TTE and markers of hemodilution^45^, maternal albumin or eGFR, but failed to identify an association between any transfer efficiencies and maternal albumin serum concentrations or estimated glomerular filtration rate. More importantly, we also observed no evidence of confounding by albumin, eGFR, or maternal weight in carefully designed longitudinal g-formula sensitivity analyses incorporating repeated maternal PFAS and these time-varying confounders.

The heterogeneity in trimester- and biomatrix-specific associations observed here between maternal serum, placental, and cord plasma PFAS levels throughout pregnancy are likely due to complex dynamics involving passive transport, PFAS-protein binding^46–48^, active transport^49–51^, gestational changes in placental development and maternal physiology, and deposition and accumulation in maternal organs such as the liver and kidneys^52–54^, the placenta, fetal cord blood, and fetal tissues^40,41^.

We applied a causal inference-informed approach leveraging comprehensive characterization of maternal serum from all three trimesters of pregnancy, the placenta, and cord blood plasma. Our results suggest that for PFOS, PFNA, and PFOA, early-to-mid pregnancy timepoints, particularly second trimester, may serve as better predictors of ultimate placental and fetal exposure burden. PFOA and PFHxS were notable exceptions when it came to cord blood levels, where placental PFOA and first trimester PFHxS were the most robust predictors. Our results may highlight higher placental to cord blood transfer for PFOA specifically, since PFOA had the lowest placental retention when comparing placental to cord levels. This may be due to, compared to PFOS and some other PFAS, PFOA has a lower binding affinity for placental proteins such as fatty acid binding protein^48^ and different binding affinities to or transport by transplacental transporters such as OAT4^49,50^. Our analysis helps to clarify whether a particular sample or combination of samples serve as the best biomarkers of placental and cord blood exposure but largely reflect prevailing knowledge about PFAS accumulation in fetal tissues throughout pregnancy^40,41^. The conventional co-adjusted GLM and shrinkage and regularization models for the cord plasma outcome appeared to suffer from bias given their divergence from g-formula results, potentially stemming from co-adjustment for downstream variables and placental levels as a potential mediator or a failure to faithfully model the true time-ordered, correlated relationships between longitudinal variables.

While our primary goal was to better understand the relationship between maternal PFAS levels and the PFAS burden carried by placental tissue and cord blood plasma, it’s essential to acknowledge that there may be window-specific or developmental effects of perinatal PFAS exposure that may not be best captured in early-to-mid pregnancy. However, our findings do help elucidate the relationship between maternal exposure and target tissues such as the placenta and fetal exposure. While direct measures of PFAS burden in tissues may best capture etiological biological effects of PFAS exposure, they are driven by internal absorption, distribution, metabolism, elimination^55^, or ADME, and a latent environmental exposure profile^56^ characterized by health behaviors and sociodemographic factors^57^, occupation^58^, and local PFAS pollution^59^. Consequently, the predominantly non-Hispanic White and well-educated cohort studied here may not be generalizable to other populations with a different risk factor profile. Future research into better characterizing internal ADME, and especially PFAS transplacental transfer and target tissue deposition mechanisms^50,55^ will further inform the study of the etiological effects of PFAS. Research effectively targeting the latent environmental exposure for intervention may yield the most cost-effective and beneficial impacts on human health, especially preventing exposure earlier in pregnancy or altogether.

From an environmental health perspective, our findings suggest priority investigation of different gestational timepoints or tissues on a PFAS-specific basis. For accurate prediction of placental PFAS levels at term, additional measurements throughout gestation may generally provide better performance than only including a single sample at any time point. However, our causal inference-informed modelling approaches suggest prioritization of earlier timepoints in pregnancy between 12 and 24 weeks for most PFAS studied to achieve good predictive capacity of placental and cord blood levels and potentially greater etiological relevance to actual placental and cord blood plasma levels. Additionally, PFOA placental levels were closely related to cord blood plasma PFOA levels for importance and prediction.

## SUPPLEMENTARY TABLES

**Supplementary Table 1** Above limit-of-detection rates for the PFAS studied.

**Supplementary Table 2** Case complete placental included vs. excluded comparison.

**Supplementary Table 3** Case complete cord blood included vs. excluded comparison.

**Supplementary Table 4** Model effect estimates for placental PFAS (ng/g). Effect estimates correspond to a doubling in maternal timepoint PFAS concentration (μg/L).

**Supplementary Table 5** Stochastic intervention g-formula model effect estimates for placental PFAS (ng/g). Effect estimates correspond to an additive shift of 0.25 standard deviations on the original scale (μg/L) pooled across maternal serum measures.

**Supplementary Table 6** Model effect estimates for cord plasma PFAS (μg/L). Effect estimates correspond to a doubling in maternal timepoint PFAS concentration (μg/L) or placental PFAS concentration (ng/g).

**Supplementary Table 7** Stochastic intervention g-formula model effect estimates for cord blood serum PFAS (μg/L). Effect estimates correspond to an additive shift of 0.25 standard deviations on the original scale (μg/L or ng/g) pooled across maternal serum measures.

## SUPPLEMENTARY FIGURES

**Supplementary Figure 1.**
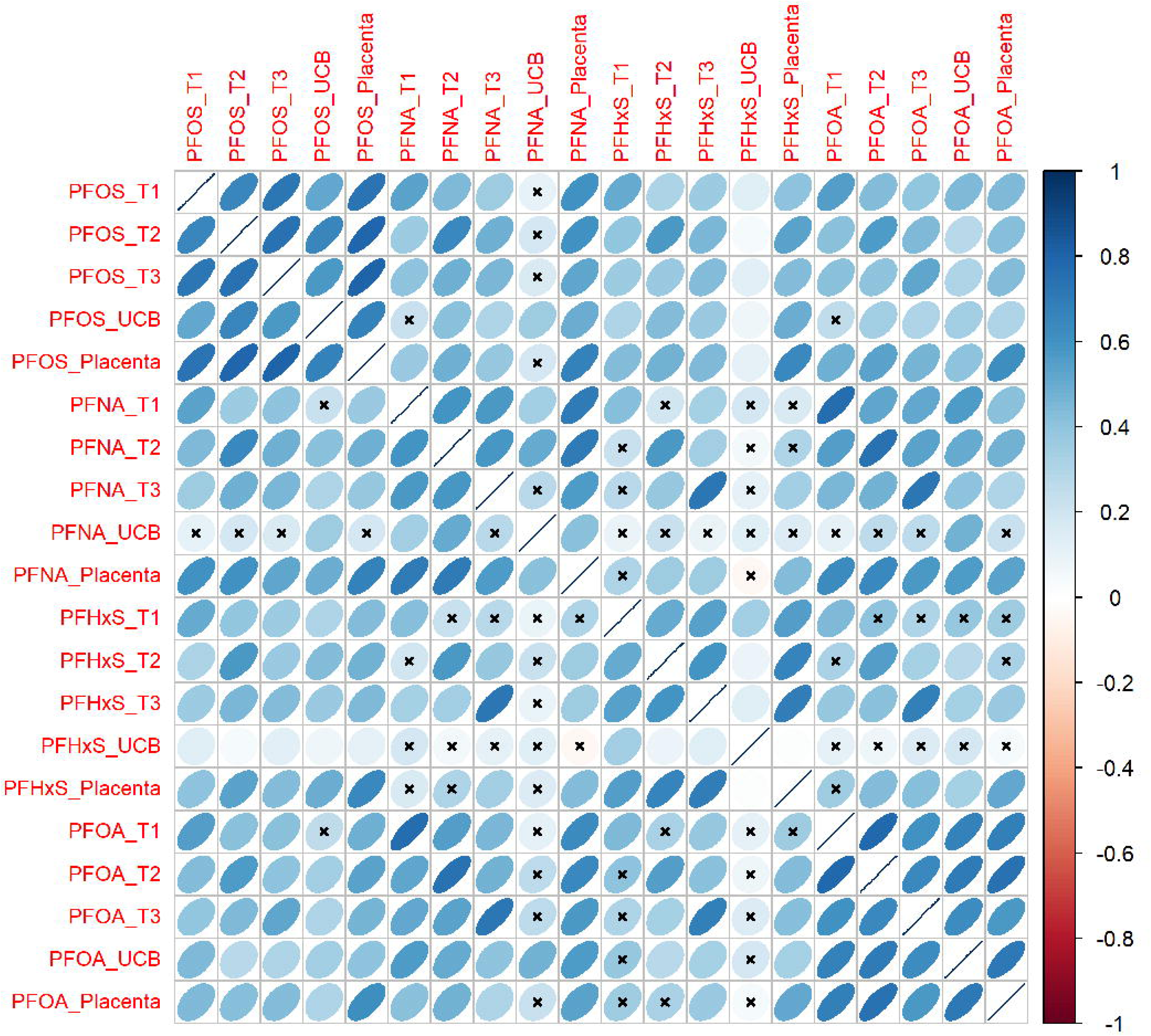
Directed acyclic graph describing the longitudinal parametric g-formula for placental PFAS as the outcome. L0 describes the vector of fixed baseline covariates: maternal body mass index, maternal age, maternal education and fetal sex. A describes maternal serum PFAS level at each timepoint (indicated with _1, _2, _3). Y describes placental PFAS level. L1 represents time-varying confounders measured at the same timepoints as maternal exposure. We tested three sensitivity models that considered L1 albumin serum level, estimated glomerular filtration rate, or maternal weight.

**Supplementary Figure 2.**
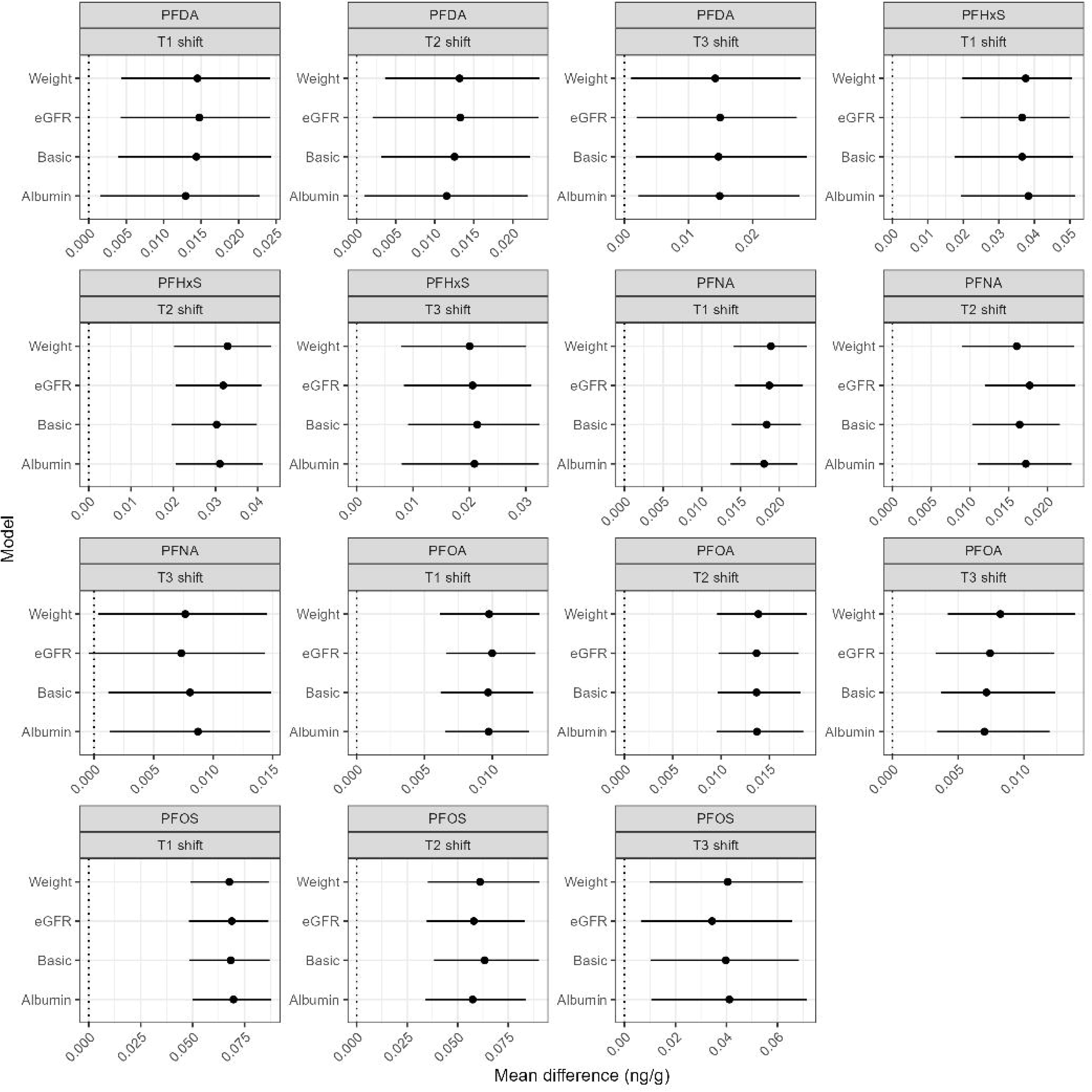
Directed acyclic graph describing the longitudinal parametric g-formula for cord blood serum as the outcome. L0 describes the vector of fixed baseline covariates: maternal body mass index, maternal age, maternal education and fetal sex. A describes maternal serum PFAS level at each timepoint (indicated with _1, _2, _3). M describes placental PFAS level. Y describes cord blood serum PFAS level. L1 represents time-varying confounders measured at the same timepoints as maternal exposure. We tested three sensitivity models that considered L1 albumin serum level, estimated glomerular filtration rate, or maternal weight.

**Supplementary Figure 3.**
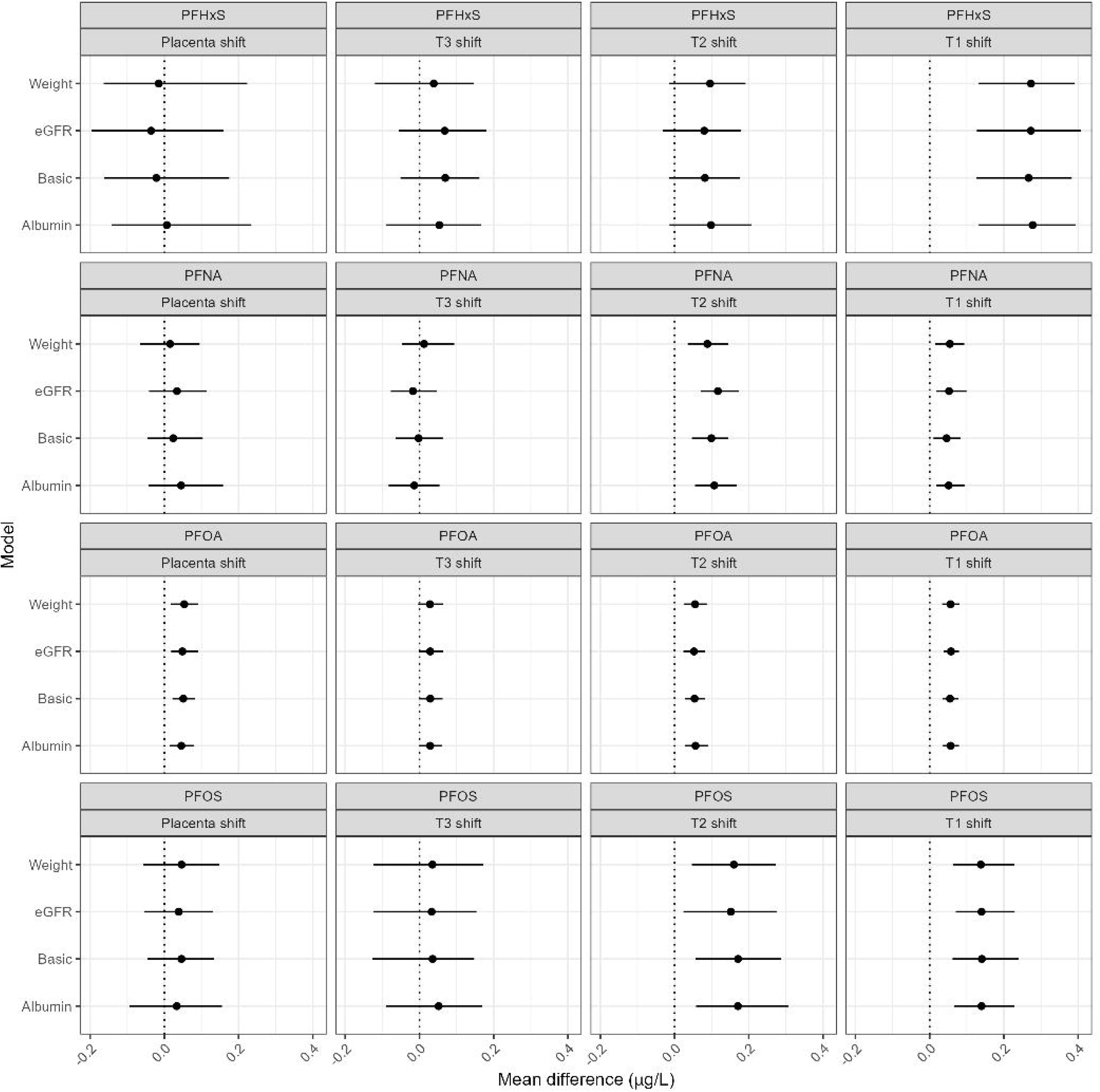
Spearman correlation matrix comparing PFAS measures across biomatrices and timepoints. An X indicates no statistically significant bivariate correlation (p<0.05).

**Supplementary Figure 4.**
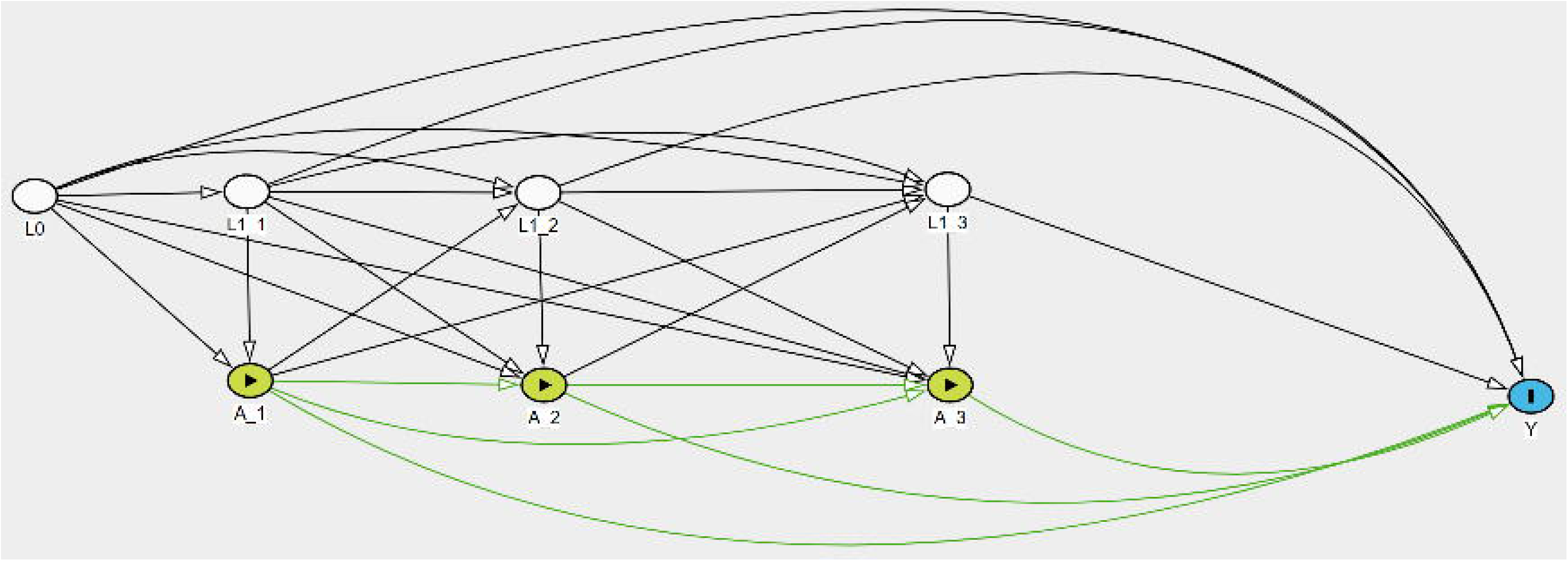
Stochastic interventions estimating the mean difference in placental concentration corresponding to a 25% standard deviation increase (shift) at one timepoint during pregnancy via the parametric g-formula with time-varying confounders albumin, estimated glomerular filtration rate (eGFR), maternal weight, or none (Basic).

**Supplementary Figure 5.**
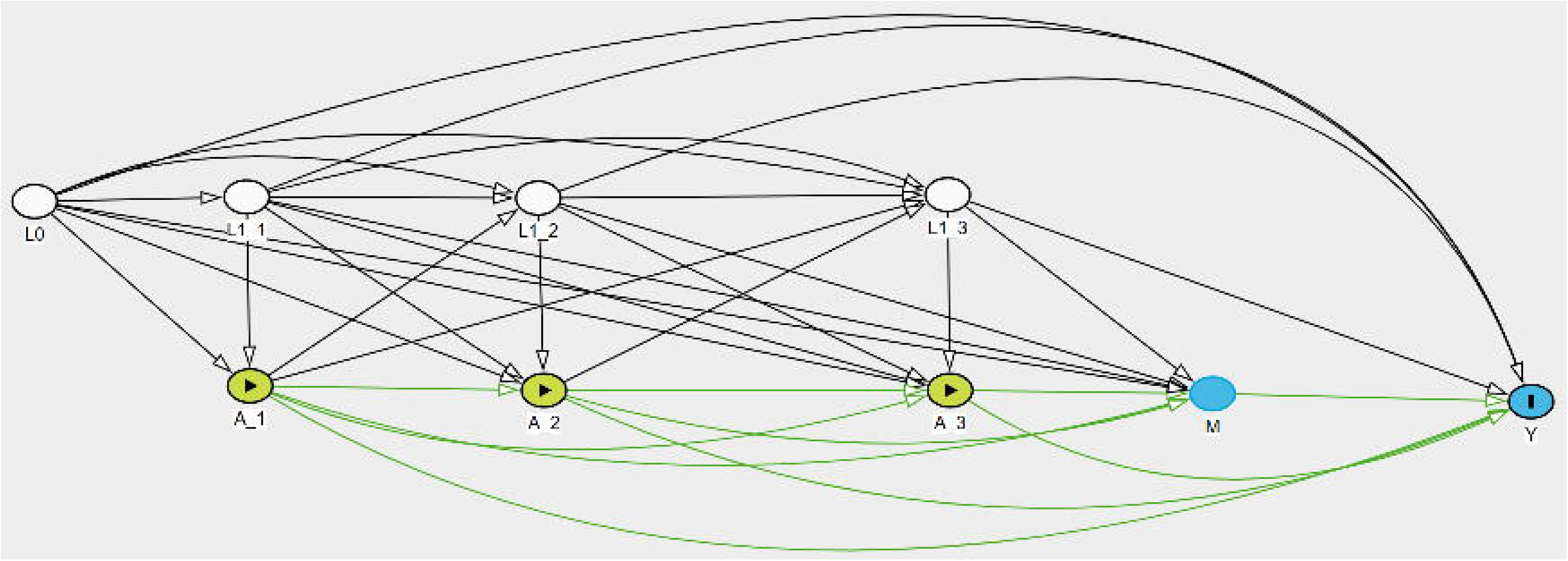
Stochastic interventions estimating the mean difference in cord blood serum concentration corresponding to a 25% standard deviation increase (shift) at one timepoint during pregnancy via the parametric g-formula with time-varying confounders albumin, estimated glomerular filtration rate (eGFR), maternal weight, or none (Basic).

## Declarations

### Data availability statement

Data are available upon reasonable request contingent on data privacy and access regulations.

## Acknowledgements

We thank the participants who provided biospecimens for this study. This work was supported by funding from the National Institute of Environmental Health Sciences, NIH/NIEHS [R01 ES032176; MPIs: Andres, Everson, & Pearson], NIH/NIEHS [R01 ES036986; MPIs: Andres & Everson], the HERCULES Center [P30 ES019776; MPIs: Everson, Barr], the UK-CARES Center [P30 ES026529; CoI: Pearson]. KS is supported by USDA ARS CRIS 3093-10700-001-000-D. The content is solely the responsibility of the authors and does not necessarily represent the official views of the National Institutes of Health. This manuscript is the result of funding in whole or in part by the National Institutes of Health (NIH). It is subject to the NIH Public Access Policy. Through acceptance of this federal funding, NIH has been given a right to make this manuscript publicly available in PubMed Central upon the Official Date of Publication, as defined by NIH.

## Author contributions statement

**Kyle Campbell:** Conceptualization, Formal analysis, Methodology, Investigation, Data Curation, Writing – Original Draft, Software, Validation, Visualization

**Dana Boyd Barr:** Conceptualization, Funding acquisition, Project administration, Resources, Supervision, Validation, Writing – review & editing

**Andrew Morris:** Data curation, Funding acquisition, Investigation, Project administration, Resources, Supervision, Validation, Writing – review & editing

**Volha Yakimavets:** Data curation, Investigation

**Parinya Panuwet:** Data curation, Investigation

**Donald Turner:** Data curation, Investigation

**Lauren A. Havens:** Data curation, Investigation

**Stephanie Eick:** Conceptualization, Methodology, Writing – review & editing

**Kartik Shankar:** Conceptualization, Data curation, Funding acquisition, Investigation, Project administration, Resources, Validation, Writing – review & editing

**Kevin J. Pearson:** Funding acquisition, Project administration, Resources, Writing – review & editing

**Aline Andres:** Conceptualization, Data curation, Funding acquisition, Investigation, Methodology, Project administration, Resources, Supervision, Validation, Writing – review & editing

**Todd M. Everson:** Conceptualization, Data curation, Formal analysis, Funding acquisition, Methodology, Project administration, Resources, Software, Supervision, Validation, Writing – original draft, Writing – review & editing

